# Posterior language areas share electrophysiological signatures of word retrieval in context-driven object and action naming

**DOI:** 10.64898/2026.05.01.721957

**Authors:** Irina Chupina, Vitória Piai, Britta U. Westner

**Author notes:** **Corresponding author:** Irina Chupina Radboud University Thomas van Aquinostraat 4 6525 GD Nijmegen, the Netherlands. These two authors contributed equally (Vitória Piai, Britta U. Westner). All supplemental materials and stimuli can be found at https://osf.io/frmh9.

## Abstract

Claims about shared neural processing between object and action words have mainly been based on spatial overlap. Spatial overlap alone, however, provides an incomplete understanding of neural (dis)similarity. Here, we compared object and action word retrieval within participants utilising temporal, spectral, and spatial information in the electroencephalogram (EEG) recorded during context-driven object and action picture naming. Constrained sentence contexts elicited pre-picture lexical-semantic word planning for object and action words, indexed by power decreases in the alpha-beta frequency range (8 – 30 Hz). Using a novel approach based on mutual information and source-reconstructed EEG signal, we computed joint temporo-spectro-spatial (dis)similarity indices across object and action naming in the constrained condition where information retrieval occurred. Spatially, dissimilarities were found in bilateral frontal, anterior superior temporal, and right anterior-to-middle temporal areas. Similarity, by contrast, was linked to the precunei and right temporo-parietal areas, regions associated with lexical-semantic processing and word retrieval. Crucially, similarity in the precunei compared to the temporo-parietal regions was characterised by differential patterns of the alpha-beta activity, implying processing and, potentially, functional differences between the areas. This finding highlights how conclusions about shared neural processes depend on the degree of abstraction (e.g., spatial, spatial-spectral) chosen to define the compared neural mechanisms. We tentatively interpret the contribution of the right hemisphere and left frontal areas to (dis)similarity as coarser, less fine-grained lexical-semantic computations.

## 1. INTRODUCTION

*Objects* and *actions* are fundamental semantic categories that are salient to infants already before they start learning their first words (Gogate & Hollich, 2016). Neural correlates of object and action word processing have been extensively studied but the extent to which word access and retrieval across the two categories is neurally similar remains under debate (for review, see Crepaldi et al., 2011, 2013; Vigliocco et al., 2011).

Comparing object and action words (labelled grammatically as nouns and verbs) is not straightforward due to inherent differences between the two categories. Multiple semantic attributes (e.g., sensorimotor attributes, imageability, argument structure) as well as morphosyntactic attributes (e.g., thematic roles, inflection) separating them cannot be entirely controlled for: Behavioural differences in response times (RTs) in picture naming persist even after matching the stimuli, suggesting that verb representations remain more conceptually, lexically and morphologically complex (Kauschke & von Frankenberg, 2008; Szekely et al., 2005). This is further reflected in the neural signatures of object and action processing, with studies demonstrating that the differences between the categories become less pronounced when semantic- and morphological-level differences, in particular, are accounted for (e.g., Elli et al., 2019; Siri et al., 2008; Vigliocco et al., 2011). When the stimuli are well-matched, lesion-symptom mapping suggests overlap between object and action naming in the left temporal and parietal regions (Alyahya et al., 2018).

On top of these intrinsic features, differences between the neural correlates of object and action word retrieval can be further exacerbated by experimental design (for more detailed discussion on the inherent and methodological differences between nouns and verbs see Crepaldi et al., 2011, 2013; Druks, 2002; Liljeström et al., 2008; Mätzig et al., 2009; Vigliocco et al., 2011). One such example is a classical visually guided word retrieval task – picture naming. It is widely used in basic research and clinical contexts (e.g., in awake brain surgery, De Witte & Mariën, 2013, and for assessment of language deficits in aphasia, Breining et al., 2022), also being the basis for findings of dissociation between noun and verb naming ability in people with neural damage due to stroke or glioma (e.g., Miceli et al., 1984; Tomasino et al., 2019). The disadvantage of picture naming when comparing objects and actions is that, even after controlling for the visual complexity of the pictures, action pictures remain more conceptually (psychologically) complex: They require more detailed “scene parsing” which involves simultaneous activation of depicted object concepts for interpreting the action (Kauschke & von Frankenberg, 2008; Szekely et al., 2005). The visual stimulus properties and their perception were reported to impact both RTs (Kauschke & von Frankenberg, 2008) and neural signatures (from electrophysiology, Amoruso et al., 2021; Fargier & Laganaro, 2015; Liljeström et al., 2008). Conversely, when object and action naming were compared using a single set of images (Fargier & Laganaro, 2015; Hernandez et al., 2001; Liljeström et al., 2008, 2009; Sörös et al., 2003), both haemodynamic measures recorded with functional magnetic resonance imaging (fMRI) and electrophysiological measures recorded with magnetoencephalography (MEG) indicated little to no spatial and temporal differences between the two categories.

To avoid visual confounds when comparing object and action word retrieval, auditory paradigms such as sentence completion can be employed. Relatively few studies, however, have elicited word production using sentential context. Abel et al. (2015) used a sentence completion task with semantically constraining sentences to analyse noun and verb production errors in healthy speakers. They observed no increased difficulty for verbs in sentence completion, but the patterns of non-target responses suggested different organisation for nouns and verbs in the mental lexicon (also see Mätzig et al., 2009). To investigate naming performance of participants with aphasia who had a selective verb deficit on picture naming, Crepaldi and colleagues (2006) used open-ended sentence completion where the target noun or verb were prompted by their derivatives (e.g., *explosion* as a prompt for the completion *explode*, in Italian). Interestingly, out of 16 participants, only two continued to show the noun-verb dissociation in this task. To the best of our knowledge, there have been no neuroimaging studies that compared auditory sentence-driven object and action word retrieval.

Importantly, the discussion so far regarding the (dis)similarities between object and action word processing has been largely based on the findings in the *spatial* domain, i.e., the overlap of activity in the brain. Accordingly, the spatial overlap of neural correlates is typically interpreted as the processing of both word categories being supported by the completely or partially overlapping underlying neural substrates. While, unarguably, spatial overlap of brain activity represents an important feature of neural similarity, it is debatable whether this observation alone provides enough ground for concluding (dis)similarities between object and action word processing (for discussion, Francken et al., 2022; Piai et al., 2025). For instance, according to Craver’s (2007) view of neural mechanisms, the spatial *and* temporal organisation of the mechanism’s components are important as changing either spatial *or* temporal aspects of the mechanism’s organisation would likely yield a different mechanism. Taking this perspective, we here argue that ultimate claims about the similarity between object and action word processing should not be made without taking other aspects of neuronal activity beyond spatial overlap into account.

### Present study

The aim of the present study was to investigate the (dis)similarities between object and action word retrieval, specifically, at the lexical-semantic planning stage, while using a more conservative criterion for what would count as similarity, that is, spectral, temporal, and spatial overlap. To this end, we obtained electroencephalography (EEG) data using an established context-driven object naming task (Griffin & Bock, 1998; Piai et al., 2014) and an analogous context-driven action naming task (see Supplement 1, https://osf.io/frmh9). In context-driven naming, participants hear either semantically constraining or unconstraining incomplete sentences that end with a picture to be named. The contextually constraining sentences have high cloze probability (Taylor, 1953) and elicit planning of the target picture name before the picture appears on screen (Chupina et al., 2025; Piai et al., 2014, 2020). Conversely, the unconstrained condition is similar to classical bare picture naming in the sense that the target word can only be planned upon picture presentation (Chupina et al., 2025; Roos et al., 2024). Therefore, contrasting the constrained to the unconstrained condition in the pre-picture interval yields an electrophysiological signature that is associated with early planning. This early planning has been shown to mainly reflect lexical-semantic processes (Piai et al., 2020) and is indexed by power decreases in the alpha (8 – 12 Hz) and beta (13 – 30 Hz) frequency bands. Spatially, the power decreases in object naming have been linked to left posterior temporal and inferior parietal areas as well as, to a lesser extent, left inferior frontal areas (e.g., Piai et al., 2014, 2018; Roos & Piai, 2020 for evidence from MEG). Importantly, although participants name pictures, the information retrieval measured before picture onset is driven by the sentential context, i.e., activation via lexical-semantic networks, mostly driven by thematic (associative) relations (e.g., *priest*, *pray* activating “*church”* or *priest*, *kneel*, *church* activating “*pray”*). This task design removes the confound of visual and conceptual picture complexity from the across-task comparison and, simultaneously, elicits word retrieval in a more naturalistic speech-production setting.

First, we designed a new context-driven action task and tested its validity (Supplement 1). Based on the cloze probabilities (Bloom & Fischler, 1980; Taylor, 1953) that were collected online for the new sentences eliciting verb production, we selected a set of best-performing stimuli and ran a behavioural experiment to establish the presence of the context effect in RTs for action naming. Following a strong and reliable behavioural context effect, we then conducted the main EEG experiment where each participant completed context-driven object and action naming tasks. On these data, we performed time-frequency analyses and source reconstruction within task as well as a between-task similarity analysis based on mutual information and the source-reconstructed EEG signal (Westner et al., 2026). Behaviourally, we expected to see the context effect within task as shorter RTs in the constrained compared to the unconstrained condition. In the EEG data, we expected the neural correlates of retrieval to be indexed by power decreases in the alpha-beta (8 – 30 Hz) frequency range in the pre-picture interval in each task, indicating lexical-semantic preplanning. Spatially, we expected to see the sources of the decreases peaking in the posterior temporal and inferior parietal regions, in line with previous findings for the object task (Roos & Piai, 2020). In terms of similarity across tasks, we hypothesised that, unlike conceptual representations or morphosyntactic processes that seem to separate objects and actions (e.g., Siri et al., 2008; Vigliocco et al., 2011), lexical-semantic retrieval might rely on shared mechanisms for both word categories (e.g., Fargier & Laganaro, 2015; Faroqi-Shah et al., 2018; Sörös et al., 2003) and, consequently, produce comparable spectral-temporal signatures in the overlapping neural regions. As we defined similarity in terms of *time-resolved spectral-spatial overlap*, all of the following conditions should be met for concluding similarity between neural signatures across the tasks: (1) the effect comes from the temporal windows where the same cognitive processes are assumed in both tasks, i.e., an interval preceding picture onset when lexical-semantic preplanning occurs; (2) the effect occurs in the same frequency bands, i.e., power modulations in the alpha and/or beta bands; (3) the effect is linked to the same cortical structures as labelled by a brain atlas.

## 2. METHODS

The protocol was approved by the Ethics Committee of the Faculty of Social Sciences at Radboud University, following the Declaration of Helsinki. Participants gave written informed consent and received monetary compensation or course credits for participation. The data were collected at the Donders Institute in Nijmegen, the Netherlands.

### Participants

Twenty-eight participants performed the auditory version of the context-driven object and action naming tasks while EEG was recorded. Two participants were excluded due to technical issues, resulting in 26 native Dutch speakers in the final sample (seven male, mean age = 21.9, SD = 2.9, range 18 – 29). Participants visited the Donders Institute twice (visit 1: EEG, visit 2: structural MRI) or once (EEG) if they already had a T1-weighted scan.

### Materials

Following validation (Supplement 1), the action-naming task set comprised 54 sentence pairs, with 25 target pictures from Druks & Masterson (2000, normed by Shao et al., 2014) and 29 pictures from VANPOP (Ohlerth et al., 2020). For the context-driven object naming, 57 grey-scale noun pictures were selected from MULTIPIC (Duñabeitia et al., 2018). Additional 23 pictures were found online. This resulted in 108 sentences (54 per condition) for the action naming task and 160 sentences (80 per condition) in the object naming task. The sentence stimuli can be found at https://osf.io/frmh9.

The target pictures with available norms were compared for visual complexity and name agreement (calculated as *h index*) across the sets. For 57 out of 80 nouns, visual complexity norms and *h indices* were available from MULTIPIC (Duñabeitia et al., 2018). For 25 verbs from Druks & Masterson’s set (2000), both measures were available from Shao et al. (2014). The remaining 29 verbs from VANPOP (Ohlerth et al., 2020) had no visual complexity norms, and we calculated their *h index* using the formula by Snodgrass and Vanderwart (1980). To compare visual complexity and name agreement across tasks, we used unpaired t-tests with Welch correction to account for unequal sample sizes and variances. Action pictures (mean = 3.7, SD = 0.77) were more visually complex than noun pictures (mean = 2.5, SD = 0.6, with t(37) = 6.7, p-value <0.001) as expected based on prior literature (e.g., Kauschke & von Frankenberg, 2008). With regards to name agreement, there were no significant differences between the noun and verb sets (t(108)=0.1, p-value = 0.92), with the mean *h index* of 0.33 (SD = 0.43) for nouns and 0.32 (SD = 0.43) for verbs, as well as between the subsets of verbs from Druks & Masterson (mean 0.35) and VANPOP (mean 0.29, with t(47) = 0.5, p-value = 0.62).

Lead-in sentences were recorded by a female native Dutch speaker at a natural speaking rate. The recordings were denoised using Audacity® (Audacity Team, 2014) and the intensity was normalised to 60 dB using PRAAT (Boersma & Weenink, 2013). Finally, the recordings were resampled to 44100 Hz and converted into stereo. The mean duration of the noun lead-in sentences was 2038 ms (SD = 310 ms, range 1391 - 3338 ms) and 2977 ms for the verb lead-in sentences (SD = 660 ms, range 1827 - 5197 ms). Each pronoun (*hij/zij*) introducing the picture sentence in the action task lasted 540 ms.

### Design and data acquisition

The experiment was programmed in Presentation (*Neurobehavioral Systems* Inc.). The presentation of stimuli was pseudorandomised using Mix (van Casteren & Davis, 2006), with a maximum of five successive repetitions of the same condition and with a minimum distance of 20 trials separating appearance of the same target picture in the two conditions. Each participant received a unique pseudorandomised list. Before the experiment, participants were familiarised with the target pictures. Participants were instructed to listen to the sentences and complete them by naming the depicted object or action. To minimise artifacts, participants were additionally instructed to stay relaxed, try not to move their jaw or head and keep the gaze at the fixation cross. EEG data were recorded using the BrainVision Recorder with the BrainAmps DC amplifier and customised actiCAP with 64 active electrodes placed according to 10-20 convention. The data were acquired at a sampling rate of 500 Hz (0.016-100Hz online band-pass filtered) with AFz as the ground electrode and TP9 placed on the left mastoid for online reference. The impedance was kept below 10 kΩ for the cap electrodes and below 5 kΩ for the ground and reference. Two pairs of external electrodes were used to record vertical and horizontal electro-oculograms, placed above and below the left eye and next to the canthi of both eyes. Another pair of electrodes recorded the electromyogram from the orbicularis oris muscle, with the electrodes placed above and below the right corner of the mouth. A 3D scan of each participant’s head with the EEG cap was made using the structure sensor by Occipital (https://structure.io/, Homölle & Oostenveld, 2019) to record the electrode positions for source reconstruction of the scalp-level activity (Dalal et al., 2014). Finally, structural T1-weighted MRI scans were acquired in a Siemens Magnetom Skyra 3T MRI scanner (Erlangen, Germany).

The order in which participants completed object and action tasks was counterbalanced. After seeing the fixation cross (1500 ms), participants heard lead-in sentences via the speakers (Figure 1). In the action naming task, the sentence was followed by a 200-ms silence and the picture sentence (*He* / *she …*). Then, in both tasks, participants saw a fixation cross for 800 ms, followed by the target picture (2000 – 2250 ms, jittered). Each trial concluded with a screen with asterisks (2000 ms). Responses were recorded with a microphone for 3000 ms, starting with picture presentation.

**Figure 1.**
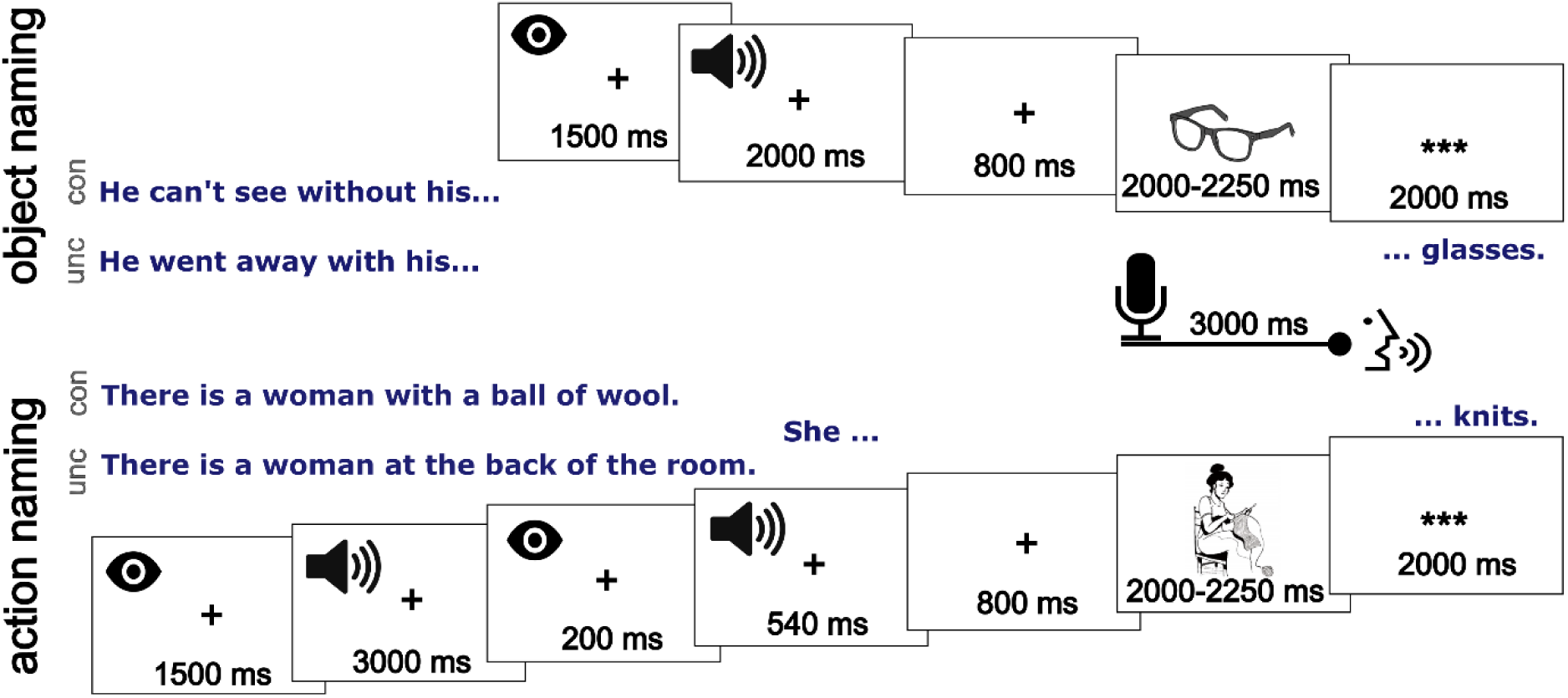
Trial design of the action and object naming tasks used in the EEG experiment. After the fixation cross, participants heard a constrained (con) or an unconstrained (unc) sentence leading to the target picture. The duration indicated for the auditory sentences represents the mean lead-in sentence duration for each task, with pronouns hij / zij (he / she) further taking 540 ms in the action task. Presentation time of the picture was jittered between 2000 and 2250 ms.

### Behavioural data analysis

RTs were calculated manually using PRAAT (Boersma & Weenink, 2013). Erroneous responses were classified as (1) general errors (omissions, word does not fit the picture, several responses in a row), (2) technical errors (correct but given before picture presentation or with interference such as coughing), (3) dysfluencies. The following responses were considered correct: (1) target words, (2) semantically close responses that fit the lead-in sentence, e.g. *schilderen* (paint) and *tekenen* (draw) for the picture *tekenen*, (3) in the unconstrained condition, words that adequately described the picture (e.g., cook instead of cut in response to the picture of a person cutting onions).

Analyses were performed in R Statistical Software (v4.1.3, R Core Team 2022). Since it is well known that action words require more time for production compared to object words (e.g., Kauschke & von Frankenberg, 2008; Szekely et al., 2005) and the constrained condition produces shorter RTs compared to the unconstrained condition (e.g., Piai et al., 2014; for action task see Supplement 1), we considered it to be more pertinent for our goal of comparing the neural activity across the tasks to focus on the RT distributions that reflect underlying cognitive processing. To illustrate RT differences between the object and the action task, we descriptively report the performance on both tasks across conditions as well as the associated effects sizes (ESs) with confidence intervals (CIs). The ESs were calculated using *unbiased Cohen’s d* for repeated design (Cumming, 2013, see Supplement 1 for more details), with 95% CIs computed using the R package MBESS (Kelley et al., 2018).

Furthermore, we conducted an ex-Gaussian decomposition analysis (e.g., Balota & Yap, 2011) to separate the bulk of the RT distributions (represented by *mu* and *sigma*, corresponding to the Gaussian mean and SD, respectively) from the tail (the *tau* parameter). The ex-Gaussian parameters were estimated per individual participant on data quantiles within condition and task using the Quantile Maximum Likelihood Estimation (Brown & Heathcote, 2003). To investigate (dis)similarities between the RT distributions, we plotted them and ran dependent samples Welch’s t-tests comparing *mu*, *sigma* and *tau* parameters per participant in each condition across the tasks.

### EEG data preprocessing and analyses

EEG data were preprocessed and analysed in MATLAB 2022a using Fieldtrip (version 20220707, Oostenveld et al., 2011). For each task, the data were segmented into epochs from 2000 ms before to 1500 ms after the picture onset. Then, baseline correction using the mean of the pre-picture interval was applied. After removing incorrect responses, each trial was manually inspected for artifacts. To remove eye-related activity, we ran an independent component analysis, discarding any components related to eye movements (on average two components per participant). Afterwards, the trials were inspected manually again to remove any remaining trials with artifacts. Finally, the cleaned data were re-referenced to the average of all channels. Trials were then separated into the constrained and unconstrained conditions.

### Within-task analyses

*Scalp level.* Using a sliding time window with a length equivalent to four cycles of the given frequency, we calculated time-frequency representations (TFRs) from 4 to 40 Hz with a 1 Hz frequency resolution. The sliding window advanced in 10 ms steps. In each time window, before performing the Fourier transform, the data were multiplied with a Hanning taper. The TFRs were calculated per trial for each participant and then averaged per condition. For visualisation, resulting TFRs were averaged across participants. Then a contrast was calculated by subtracting the average TFR of the unconstrained from the average TFR of the constrained condition and normalising the results by the average of the two conditions. The visual inspection of the TFRs (see *Results*) suggested that the alpha-beta power decrease was shifted in the action task relative to the object task, with the action task effect occurring earlier. To better align the neural activity across the tasks (i.e., ensure that the time windows of interest reflect similar cognitive processes), we found the latency of minimal value in the alpha-beta frequency range (i.e., the maximal power decrease in microvolts) for either task and centred the 800 ms intervals on this latency. This resulted in an interval of – 800 to 0 ms for the object task and –1150 to –350 ms for the action task. All further within-and between-task analyses were conducted within these time windows. To establish if there were electrophysiological differences between constrained and unconstrained conditions within the 800 ms intervals of interest, we used non-parametric cluster-based permutation tests (Maris & Oostenveld, 2007). Per task, we conducted a dependent samples t-test at every available channel-time-frequency sample, with the samples exceeding the threshold of p < 0.05 (one-tailed) used for clustering. To form a cluster, significant samples had to be a minimum of two neighbouring channels adjacent in time and frequency, with the cluster-level statistic calculated as the sum of t-values within each cluster. This clustering procedure was repeated 1000 times using random permutations of the tested conditions. Finally, the Monte Carlo p-value of the obtained clusters was computed (alpha level of 0.05, one-tailed).

#### Source level

To reconstruct the sources for the alpha-beta power decreases in each task, we first created a realistic boundary element head model (BEM). The individual MRI scans were resliced, realigned and segmented into brain, skull and scalp using SPM12 (Version 7771, Penny et al., 2011). From the meshes for brain, skull, and skin, we then prepared a volume conduction model of the head for each participant, with the conductivity values for brain, skull and scalp of 0.33, 0.0041 and 0.33 S/m. Next, a source grid with a resolution of 10 mm was created for each subject in the Montreal Neurological Institute (MNI) space to enable comparison across participants. The positions of the EEG cap electrodes on the participants’ heads were manually marked using the 3D images from the structural scanner. Next, the electrode positions were coregistered with the MRI scans using a custom-written surface matching algorithm to align the structural scan and headmodel (Dalal et al., 2011). These coordinates together with the volume conduction model and the normalised grid were used to create the forward model.

Using the DICS beamforming approach as the inverse solution (Gross et al., 2001), we reconstructed the sources for the alpha-beta activity within the 800 ms time windows for each task within-participant. To this end, we first computed the cross-spectral density (CSD) matrix of the sensor-level data (for both conditions combined and each condition separately) at 14 Hz with a spectral smoothing of 6 Hz, to cover our frequency band of interest from 8 to 20 Hz. We computed a common spatial filter for each grid point (unit-noise-gain normalisation; Tikhonov regularisation with 5% of sensor space power). Finally, the common filter was applied to the CSD matrices computed for each condition to obtain the source power per condition. For visualisation, the difference between conditions was calculated at the subject level by subtracting the unconstrained from the constrained source power and normalising the result by the average between the conditions. Finally, the normalised differences between conditions were averaged across participants for each task.

### Between-task analyses

To investigate the electrophysiological dis(similarities) across the tasks, we conducted a similarity analysis based on mutual information in the source-level EEG signal (Westner et al., 2026). For both tasks, only the trials of the constrained condition were used since they reflect retrieval of information pre-picture. After creating an 8 mm template grid, we used the Brainnetome atlas (Fan et al., 2016) to arrange the data from grid points into parcels. Subcortical structures where EEG sources are less reliably reconstructed were manually excluded. Small adjacent parcels were merged to prevent parcels with fewer than three grid points. This yielded 126 parcels across both hemispheres (out of 246 originally in Brainnetome). Within each parcel, we used singular value decomposition to compute the strongest leadfield component. Then, the source-reconstructed activity in the time window of interest (see *Within-task analysis*) was obtained for each parcel using a linearly constrained minimum variance (LCMV) beamformer. The spatial filter was computed across both tasks (unit-noise-gain normalisation, Tikhonov regularisation with 15% of sensor space power 15% lambda for Tikhonov regularisation) to allow for their comparison. Next, these source data were used to produce the time-frequency representations at the single-trial level. Finally, the time-frequency-resolved data were log-transformed.

The similarity analysis on these data (for more detail, see Westner et al., 2026) was performed at the subject level using Python (Version 3.12.4). Briefly, we first computed a mutual information curve within task to serve as a reference for maximal similarity. For this, the data were split into two halves. From these halves, the trials were randomly subset, choosing n trials at a time and increasing n, starting from comparing the one trial to one trial and ending at comparing the averages of all trials of both halves. The operation was repeated 100 times, producing an averaged curve. Next, following the same procedure when picking sets of trials from object and action naming tasks, the computations were repeated across tasks and compared to the within-task similarity (see Supplement 2, Figure S2A-1, https://osf.io/frmh9). Then, we quantified the impact of 81 timepoints, 27 frequencies and 126 parcels on the similarity in the data. This was done by leaving one feature out and registering the impact on the mutual information by comparing the mutual information values of half of the data sets with and without this feature. Thus, to see how similar the tasks were, for instance, at 10 Hz, we left this feature of interest out, recomputed the mutual information between one half of the object and one half of the action data and then compared the value to the value when this feature was included. If the similarity decreased, the removed feature was more similar between the tasks, and vice versa. To account for variation by the random half-splits of the data, we repeated this procedure 100 times. The *similarity index* (i.e., the averaged resulting change in mutual information) was then multiplied by –1 to make the index increase for similar features and decrease for dissimilar. Using cluster permutation across participants (as implemented in MNE-Python, version 1.7.0; Larson et al., 2024; Gramfort et al., 2013), we tested against zero for the null hypothesis that leaving out each feature does not impact the similarity index for timepoints, frequencies and parcels. We assumed a linear neighbouring structure for frequencies and timepoints. For parcels, neighbours were identified based on the distance of the centres of mass among parcels.

Finally, we explored the contribution of specific frequencies and timepoints to the (dis)similarity observed in the significant parcels. If several parcels came from the same anatomical area (e.g., inferior parietal lobule), we merged them into one brain area. Taking raw source-level data in the constrained condition, we plotted them within each area (per task, averaged across participants) as a function of time and frequency and then subtracted action from object task TFRs to obtain the difference. Given that the time windows were not perfectly aligned and the end of the interval might be capturing different cognitive processes across the tasks (see *Within-task analyses*), we mainly focused on interpreting activity within the first 500 ms of the intervals.

## 3. RESULTS

### Behavioural results

The cumulative and density RT distribution plots (Figure 2) demonstrated that the constrained condition distributions across the tasks looked similar. In both tasks, the fastest responses in the constrained condition were below 450 ms. Actions, however, were named slower on average. Participants also committed more errors in the action compared to the object task (90 or 3.2% and 59 or 1.4% of total responses, respectively). Descriptive, ex-Gaussian, and effect size statistics can be found in Table 1. In the constrained condition, the bulk of the RTs (*mu* and *sigma*) was distributed without any statistically significant differences between object and action tasks, while the tails (*tau*) were statistically different. In the unconstrained condition, by contrast, both the bulk (*mu*) and the tail (*tau*) were statistically different across the tasks, with the spread of the bulk (*sigma*) not showing any differences.

**Figure 2.**
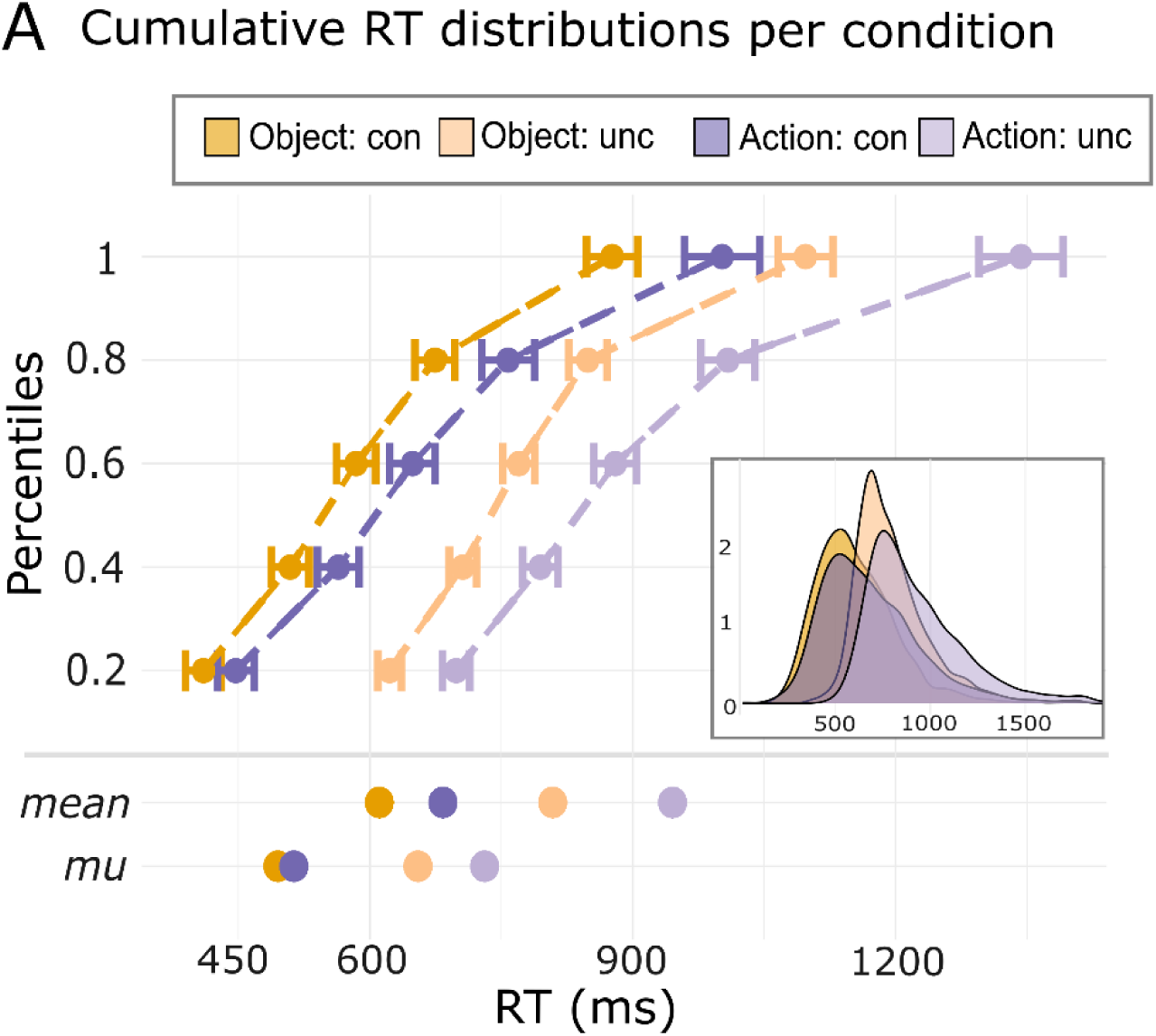
Distributions of RTs for constrained (con) and unconstrained (unc) condition for object and action naming. Cumulative RT (in quantiles) and density distributions for object (dark orange constrained, light orange unconstrained) and action (dark purple constrained, light purple unconstrained) tasks with their respective mean and *mu* values below.

**Table 1.** Descriptive, ex-Gaussian and effect size statistics for object and action tasks. All measures are in millisecond, effect sizes are measured as Cohen’s d with an adjustment (see Supplement 1 for more detail).

|  | Mean<br>(SD) | Context<br>effect<br>on<br>mean | d on<br>mean<br>[CI] | Mu | Sigma | Tau | Context<br>effect<br>on mu | d on mu<br>[CI] |
| --- | --- | --- | --- | --- | --- | --- | --- | --- |
| Object naming |  |  |  |  |  |  |  |  |
| Constrained | 611<br>(178) | 198 | 1.82<br>[1.2 –<br>2.5] | 495 | 95 | 112 | 160 | 1.36<br>[0.8 – 2] |
| Unconstrained | 809<br>(193) |  |  | 655 | 60 | 147 |  |  |
| Action naming |  |  |  |  |  |  |  |  |
| Constrained | 684<br>(209) | 261 | 1.87<br>[1.2 –<br>2.5] | 513 | 101 | 175 | 218 | 1.64 [ 1<br>– 2.3] |
| Unconstrained | 945<br>(256) |  |  | 731 | 77 | 204 |  |  |
| Dependent t-tests for ex-Gaussian parameters across tasks [action vs object naming] |  |  |  |  |  |  |  |  |
| Constrained |  |  |  | t(25)= 0.7,<br>p = 0.5,<br>CI [–33,<br>70] | t(25)= 0.4,<br>p = 0.7,<br>CI [–22,<br>34] | t(25)= 2.4,<br>**p = 0.02**,<br>CI [10,<br>118] |  |  |
| Unconstrained |  |  |  | t(25)= 5,<br>**p < 0.001**,<br>CI [44,<br>108] | t(25)= 1.2,<br>p = 0.2,<br>CI [–11,<br>44] | t(25)= 3,<br>**p < 0.01**,<br>CI [18, 96] |  |  |

Note. CI = confidence interval, SD = standard deviation.

The context effect (i.e., facilitation in the constrained compared to the unconstrained condition) was considerably larger in the action task. Participants named pictures in the constrained condition on average 198 ms faster than in the unconstrained condition in the object task (160 ms for the mu) and 261 ms faster in the action task (218 ms for the mu). The *unbiased Cohen’s d* was 1.82 for the mean in the object task and 1.87 in the action task. *Cohen’s d* for the *mu* estimate was 1.36 and 1.64, respectively. At the individual level, the magnitude of the raw context effect in the object task moderately correlated with the magnitude in the action task (r = 0.64, p < 0.001), meaning that the larger the effect was in the object task, the larger it was in the action task. Although there was some variability across participants, the context effect in the action task was typically larger than the effect in the object task.

### Within-task EEG results

We observed power decreases in the alpha-beta frequency bands (8 – 30 Hz) in the constrained as compared to the unconstrained condition in the pre-picture interval in both tasks (Figure 3A). Based on the timing of the alpha-beta decreases, participants seemed to have started target-word preplanning immediately after the constraining lead-in sentence in both tasks (Figure 3B). Thus, in the object task, decreases peaked in the 500 ms before the picture. In the action task, the activity peaked around 1000 – 500 ms before the picture, suggesting that lexical-semantic preplanning partially overlapped with the auditory pronouns *he*/*she* that introduced the picture sentence (Figure 1 and 3B). The non-parametric cluster-based permutation tests of the differences between conditions (4 – 40 Hz, within the 800-ms intervals of interest, Figure 3B) yielded negative clusters for the object task (p = 0.004), with the most prominent power decreases between –700 and 0 ms across all tested frequencies, and the action task (p = 0.002), with the most prominent power decreases between –1150 and –350 ms and 4 – 28 Hz. Source reconstruction results for the alpha-beta power decreases within-task (8 – 20 Hz, showing source points with values below a cutoff of 50% of minimum power, see Figure 3C) indicated that activity was overall bilaterally distributed. The signature of object preplanning localised to parts of bilateral temporal lobes and parietal areas, with some activity in the frontal lobes. The sources for action preplanning were also found in the bilateral posterior temporal and left parietal areas, together with more extensive activity in the frontal areas in the right hemisphere.

**Figure 3.**
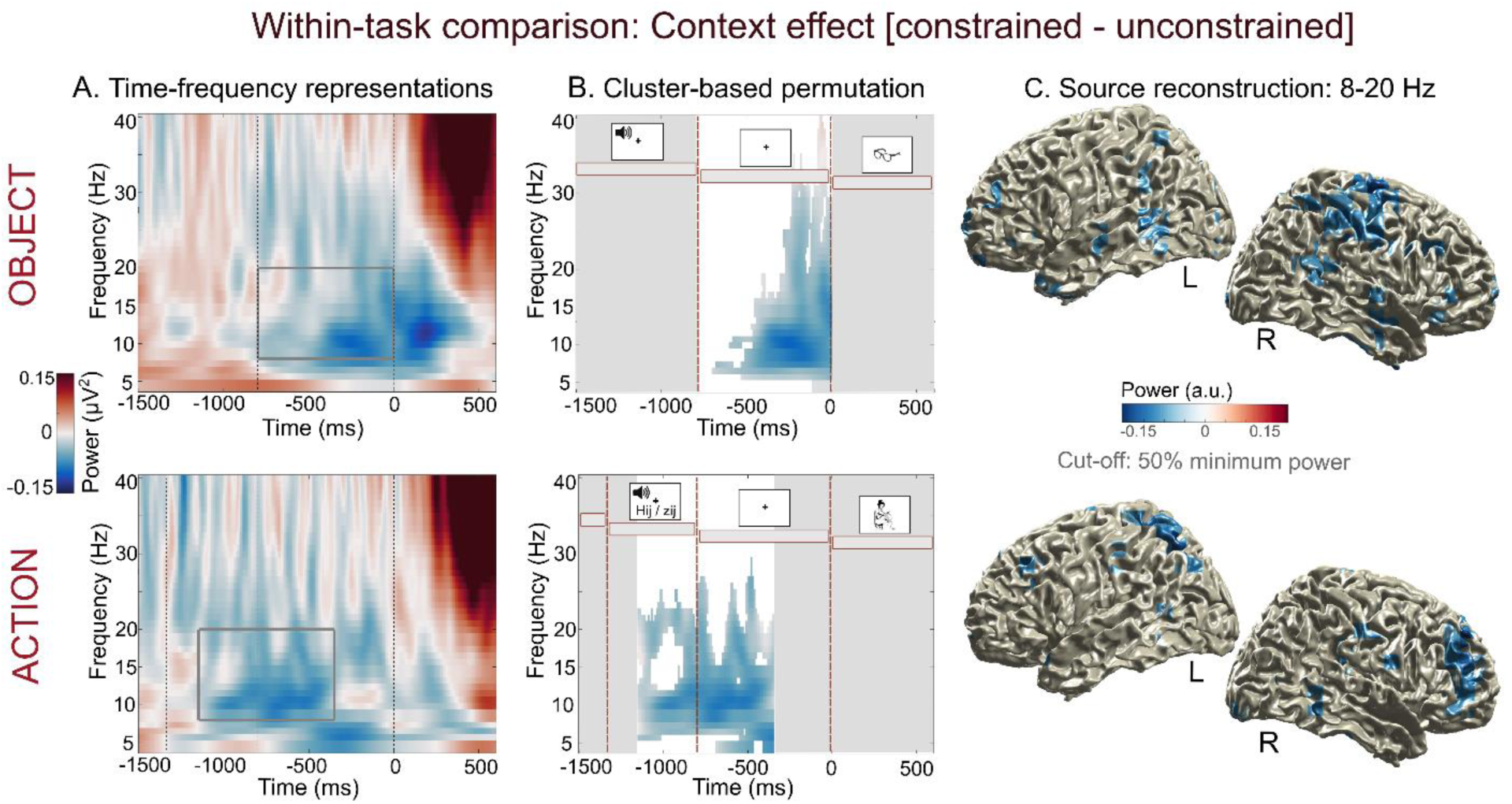
Within-task EEG results. The electrophysiological context effect (constrained – unconstrained condition, relative difference) within object (*top panels*) and action (*bottom panels*) tasks. **A**. Group-level time-frequency representations (4 – 40 Hz). The intervals containing maximal power decreases that were used for within-task source and between-task analyses are framed by grey rectangles. The stimulus events are marked by dotted lines. In both tasks, pictures were presented at zero. In the object task, the auditory lead-in sentence finished at –800 ms. In the action task, the dotted line at –800 ms marks the end of the pronoun *he / she*, while the lead-in sentence offset is marked 540 ms earlier. **B.** Cluster-based permutation results, showing differences in activity across conditions over all channels and frequencies that survived the correction according to the Monte Carlo procedure (alpha 0.05, one-sided). The dotted red lines mark sentence offset and picture presentation events. The time windows where differences were not statistically tested are marked as shaded grey areas. **C.** Brain surface plots showing the sources of the context effect (8 – 20 Hz) at the group-level in left (L) and right (R) hemispheres. The shown activity is masked at 50% of power decreases (cutoff value: 50% of minimum power).

### Between-task EEG results

The results of the similarity analyses conducted for time, frequency and parcels can be found in Figure 4. In the temporal domain (Figure 4A), there were no significantly similar or dissimilar timepoints. Descriptively, the earlier timepoints demonstrated more similarity than the later timepoints. Speculatively, this pattern might have arisen due to the shift of the analysis interval in the action task to an earlier time window, thus causing more divergence at the end of the time window that approached picture presentation in the object but not action task (see Figure 3A). Simultaneously, the activity across tasks in the earlier part of the time window might have not been sufficiently similar due to the additional presence of the auditory pronoun in the action task. In the spectral domain (Figure 4B), the activity across tasks over all parcels and timepoints was similar in the alpha band range (8 – 13 Hz, peaking around 10 Hz) and dissimilar in the beta band (17 – 30 Hz, peaking around 23 Hz).

**Figure 4.**
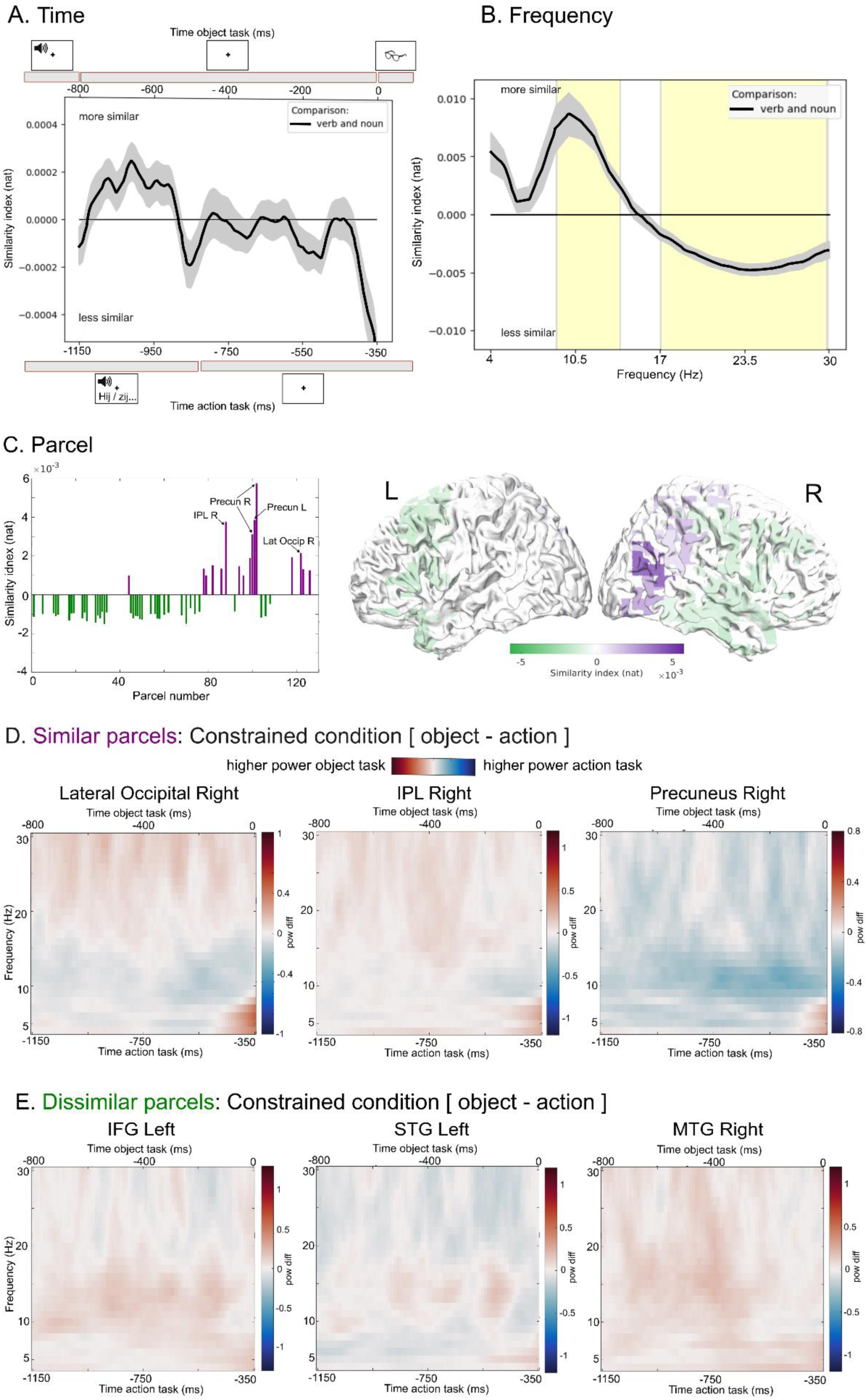
Between-task similarity analysis results. The similarity analysis was conducted on the source-level data of the constrained condition in the pre-picture interval. The higher the similarity index, the more similar the signal is (and vice versa). The impact statistics were computed using cluster-based permutation. **A**. Impact of timepoints on (dis)similarity across tasks. The x-axis at the top of the figure shows the trial event timing in the object task, and the x-axis at the bottom shows the trial event timing in the action task. There were no statistically (dis)similar timepoints (over all frequencies and parcels). **B**. Impact of frequency on (dis)similarity across tasks. The yellow shaded areas represent statistically similar and dissimilar frequencies (over all timepoints and parcels). **C**. Impact of parcel on (dis)similarity across tasks. *Left panel:* Out of 126 parcels, 16 showed similarity (purple) and 38 showed dissimilarity (green). Most similar parcels are labelled in the plot. *Right panel*: The statistically (dis)similar parcels as plotted on the cortical surface (L = left hemisphere, R = right hemisphere). **D**, **E**. Time-frequency representations of the difference between tasks (object – action) in selected similar and dissimilar parcels. The x-axes at the top and bottom of the plots show the timing in the object and action task, respectively. Values around zero represent most similarity between the tasks, positive values (red) – higher power in the object task and negative values (blue) – higher power in the action task. The colourbar limits are scaled to the limits of the time-frequency representations for object and action tasks. *Note*. Precun R = right precuneus, Precun L = left precuneus, Lat Occip R = lateral occipital right, IPL R = right inferior parietal lobule, IFG = inferior frontal gyrus, STG = superior temporal gyrus, MTG = middle temporal gyrus, ms = millisecond.

Spatially, we found 16 similar and 38 dissimilar parcels (Figure 4C, left panel) that were part of clusters in the permutation test. For interpretation, we merged parcels in clusters that were falling within the same anatomical brain area (e.g., gyri, lobules). This resulted in 8 similar and 15 dissimilar brain areas (see Supplement 2, Figure S2A-2, https://osf.io/frmh9/). The parcels showing most similarity were in the bilateral precuneus, right inferior parietal lobule (IPL) and right lateral occipital cortex, with bilateral superior parietal lobules (SPLs), left occipital regions and right precentral gyrus also showing similarity. Areas of dissimilarity were the bilateral inferior frontal gyri (IFG), middle frontal gyri (MFG), superior temporal gyri (STG), orbital, and precentral gyri, as well as left superior frontal gyrus (SFG), right middle temporal gyrus (MTG), inferior temporal gyrus (ITG), postcentral gyrus, and a subarea of right IPL. Overall, frontal and anterior-to-middle temporal regions showed dissimilarity, whereas posterior temporal and parietal areas showed similarity (Figure 4C, right panel). Interestingly, apart from the dissimilarities in the frontal and anterior superior temporal areas in the left hemisphere, most effects, including similarity, were observed in the right hemisphere.

Additionally, we plotted the power difference between the object and action task time-frequency representations within brain areas to explore the contribution of specific frequencies and timepoints (Figure 4D and 4E, plots for all brain areas can be found in Supplement 2, Figure S2A-2). After the subtraction (object – action), values around zero represent most similarity, positive values (red) represent higher power in the object task and negative values (blue) higher power in the action task. Amongst the similar parcels (Figure 4D and Figure S2A-2), IPL and SPL activity patterns can be descriptively clustered together, demonstrating most similarity in the alpha/beta frequency range. The bilateral precunei showed a different pattern, with similarity observed mainly in the beta band, with overall higher power in the action task across both bands. Finally, similarity in the bilateral lateral occipital areas was mainly driven by activity in the beta band, with slightly higher power in the action task throughout the alpha/lower beta frequency bands. The patterns of dissimilar activity in the frontal lobes and anterior STG look similar bilaterally, showing higher power in the object task in the alpha/lower beta frequency range and higher power in the action task in the beta range (Figure 4E and Figure S2A-2). A slightly different pattern emerged in the right MTG and ITG areas, where more power was observed in the object task across the whole alpha-beta frequency range (8 – 30 Hz).

## 4. DISCUSSION

We set out to investigate (dis)similarity between object and action word processing using measures capturing temporal, spectral and spatial features of neural activity. To that end, we compared electrophysiological lexical-semantic retrieval signatures between context-driven object naming (Griffin & Bock, 1998; Piai et al., 2014) and an analogous action naming task.

Within-task, the semantically constraining context of the lead-in sentences was associated with shorter RTs in both tasks, indicating preplanning of word representations before picture onset (e.g., Chupina et al., 2025; Piai et al., 2014). In line with prior literature, participants were faster at object naming compared to action naming (e.g., Kauschke & von Frankenberg, 2008; Mätzig et al., 2009; Szekely et al., 2005). While this object naming RT advantage still held throughout the whole distribution in the constrained condition, the difference between the *mus* across the tasks (i.e., the bulk of the RTs) was considerably smaller in the constrained compared to the unconstrained condition (Table 1). Together, the RT results indicated higher similarity between the two tasks in the constrained condition, where lexical-semantic preplanning occurs (Chupina et al., 2025; Piai et al., 2014; 2020) compared to the unconstrained condition, where word retrieval starts upon picture presentation. In other words, word planning elicited via sentential context seems to result in more (cognitively) similar naming behaviour. This might be tentatively explained by the fact that the confounds related to picture processing (e.g., Kauschke & von Frankenberg, 2008) were present in the visually guided (unconstrained condition) but not in the context-driven (constrained condition) word retrieval.

The difference between conditions in the pre-picture interval was indexed by power decreases in the alpha-beta frequency bands (8 – 30 Hz) (e.g., Piai et al., 2014) in both tasks. Spatially, these object and action word preplanning signatures (8 – 20 Hz) were localised to the bilateral frontal, posterior temporal, and parietal areas (Figure 3C), (partly or mostly) in line with prior MEG literature on object naming (e.g., Piai et al., 2015; Roos & Piai, 2020).

To investigate (dis)similarities between object and action word preplanning, we compared pre-picture activity in the constrained conditions. Rather than examining spatial overlap of activity in the neural structures, which is commonly used as evidence for shared processing (e.g., Faroqi-Shah et al., 2018), we jointly assessed the shared spectral, temporal and spatial information. Assuming the realignment of the pre-picture interval in the action task adequately matched the windows of lexical-semantic information retrieval across the tasks (see *Within-task analyses* and *Limitations*), the main findings were: (a) similarity was linked to posterior temporal and parietal areas while dissimilarity was linked to frontal and anterior-to-middle temporal areas, (b) differential contributions of alpha and beta band activity within brain regions formed multiple (dis)similarity patterns, with overall similarity driven by the alpha band, and (c) the majority of the (dis)similar regions was found in the right hemisphere. Below, we discuss these findings in more detail, followed by limitations and future directions.

### Shared signatures of lexical-semantic processes in the posterior but not frontal language areas

The spatial (dis)similarity patterns between the object and action task (Figure 4C) indicated a clear distinction between the more dissimilar anterior (bilateral frontal and anterior-to-middle temporal) and the more similar posterior (right posterior temporal and parietal) brain areas (for discussion of the right-hemispheric bias, see below).

Following prior literature, the dissimilarity in the frontal and anterior-to-middle temporal areas could be attributed to the inherent object-action category differences. In particular, semantic- and morphological-level differences could have contributed to the differences between the neural signatures across the tasks (e.g., Vigliocco et al., 2011). According to grounded cognition models postulating that semantic-level representations are modality-specific (Barsalou, 2008; Barsalou et al., 2003), object and action concepts would be differentially distributed in the brain due to dissimilarities in motor, visual and auditory features between them (Martin, 2016; Martin & Chao, 2001; but see Tyler et al., 2003).

Indeed, when controlling for semantic differences, Siri et al. (2008) found that picture-naming fMRI signatures for objects and actions largely overlapped except for in the left IFG, which was modulated by morphological complexity rather than the grammatical words class. Thus, we could associate the more visually salient object representations with the temporal lobe dissimilarities that we find (e.g., Wise et al., 2000). In turn, action words, which had to be inflected in the third person singular in our study, were more syntactically complex compared to object words, thus engaging IFG to a higher extent (e.g., Chang et al., 2018).

Another factor that could have contributed to the differences in the IFG activity is the extent of phonological preplanning across tasks. While the RT advantage in the constrained contexts has been primarily linked to the retrieval of lexical-semantic information of the upcoming target word (Gastaldon et al., 2020; Piai et al., 2014), under certain circumstances, the preplanning can go as far as the phonological/phonetic stage (Chupina et al., 2025). While the results of Bayesian modelling of the RTs in the present experiment were inconclusive regarding phonological preplanning (see Supplement 2B), the cumulative RT plots for longer and shorter words indicated its potential presence at least on some trials in the object task (Figure S2B-2). Given the association of phonological encoding with the IFG (de Zubicaray, 2023), participants not preplanning phonology as consistently (or at all) in the action task might explain the dissimilarity patterns in the frontal lobe.

Regarding spatial similarity, the similar parcels in the posterior temporal and parietal areas are consistent with the previously reported spatial alpha-beta signatures observed in context-driven object naming in the left hemisphere (Piai et al., 2015; Roos & Piai, 2020). These signatures have been linked to retrieval of lexical-semantic information by Piai and colleagues (2020), who demonstrated that the power decreases were sensitive to semantic information in the distractor words presented in the pre-picture interval (see also Chupina et al., 2025). Our similarity findings are also in line with MEG studies of visually guided object and action naming, which reported bilateral activity in the occipital, posterior temporal and parietal areas in the time window associated with lexical-semantic processing (approximately 200 – 400 ms, Liljeström et al., 2009; Sörös et al., 2003). Using object picture naming, Levelt and colleagues (1998) also showed sources in the right parietal cortex around 150 – 275 ms, linking them to lemma selection.

Our results based on the spatial information in the EEG signal are also somewhat consistent with previous MRI-based studies that reported spatial overlap between object and action processing (see also next section). Areas of similarity from our study partly overlap with the homologous areas (i.e., anatomic equivalents in the other hemisphere) in the left temporo-parietal regions highlighted as shared in object and action naming by a lesion-symptom mapping study by Alyahya and colleagues (2018). Furthermore, they overlap with another homologous area in the left lateral inferior temporal lobe (part of BA37) reported as shared across the tasks by Faroqi-Shah and colleagues (2018) in their activation likelihood estimation meta-analysis based on fMRI and positron emission tomography (PET) studies. The authors linked this area to modality-independent single-word lexical access. Finally, according to Binder and colleagues (2009), the bilateral angular gyri (forming part of the IPL) play an important role in semantic processing, in particular, conceptual retrieval and integration in sentence comprehension and planning, both of which are involved in context-driven naming.

### Alpha-beta activity differentially contributes to spatial (dis)similarity across brain areas

We found the spectral fingerprints of word retrieval in the object and the action tasks to be overall similar in the alpha and lower beta range (9 – 14 Hz) and dissimilar in the higher beta frequency band (17 – 30 Hz) (Figure 4B). Speculatively, the overall similarity mainly in the alpha but not higher beta band might be explained by the dominance of the alpha rhythm in neural computations (for functional interpretations, see below). Being ubiquitous throughout the brain, the alpha rhythm seems to subserve an array of cognitive functions via multiple neural mechanisms (e.g., Clayton et al., 2018; Pascucci et al., 2025). Thus, its presence in multiple parcels across the tasks might index various (un)related processes, all of which are supported by computations in the alpha frequency.

Concerning spectral signatures within individual brain areas, we observed differential alpha- and beta-band power modulation patterns across similar and dissimilar regions (Figure 4E, 4D and Figure S2A-2). Amongst the similar parcels, right IPL and SPL areas demonstrated a similar signature across tasks in the alpha and beta band. This location corresponds to homologous areas in the left hemisphere where context-driven decreases are consistently reported (see previous section). Importantly, the alpha-beta decreases specifically in these regions have been found to be critical for the presence of the behavioural facilitation in the constrained condition (Piai et al., 2018). As demonstrated by the similarity and within-task signatures, power changes in the alpha and beta bands co-occurred in our task (also see Piai et al., 2014; Roos & Piai, 2020), similarly to findings in the sensorimotor (e.g., Pfurtscheller, 1981) and long-term memory domains (e.g., Griffiths et al., 2021; for language, also see Zioga et al., 2023). While it is unclear whether the two bands together index one or a combination of processes, their strong temporal co-dependence and the finding that the decreases in the two bands in context-driven object naming are spatially indissociable in the left posterior temporal and inferior parietal but not frontal regions (Cao et al., 2022) suggest that this particular alpha-beta activity could be interpreted as one signature (but see Westner et al., 2026 for discussion on language vs episodic memory).

In the language literature, alpha and beta oscillations have been reported both in production and comprehension (for review see Meyer, 2018; Piai & Zheng, 2019; Prystauka & Lewis, 2019), with decreases linked to lexical-semantic processes (e.g., Bastiaansen et al., 2005; Piai et al., 2020). For the functional interpretation, authors have typically adopted accounts from perceptual and memory domains (e.g., Piai & Zheng, 2019; Zioga et al., 2024). Following the theory of Hanslmayr and colleagues (2012), the alpha-beta power decreases might indicate retrieval of lexical-semantic information, with larger decreases indexing higher informational content. Alternatively, the co-occurring alpha-beta activity might reflect disinhibition (i.e., decreases) in the alpha band signalling higher degrees of cortical activation in the task-relevant areas (Jensen & Mazaheri, 2010; Klimesch, 2012; Klimesch et al., 2007) coordinated with maintenance and/or (re)activation of task-specific information associated with the beta band (Engel & Fries, 2010; Spitzer & Haegens, 2017). Interestingly, the fact that alpha-beta power decreases correlate negatively with BOLD increases (Conner et al., 2011; Hanslmayr et al., 2011; Ritter et al., 2009; Scheeringa et al., 2009, 2011) makes these functional spectral-spatial interpretations consistent with the fMRI findings where spatial overlap in the homologous left-hemispheric areas was similarly interpreted as indicative of shared lexical access in noun and verb processing (Faroqi-Shah et al., 2018). Finally, although prior empirical findings support the lexical-semantic nature of the context-driven alpha-beta power decreases (e.g., Piai et al., 2014; 2020), the pre-picture activity might have also been (partly) driven by the oculomotor activity that has been implicated in encoding and retrieval in the memory domain and likely extends beyond it (Popov & Staudigl, 2023).

In the bilateral precuneus, similarity arose in the beta band while overall power was higher in the action task throughout the whole alpha-beta range. The precunei are well-connected to a widespread network of higher-association (sub)cortical structures, including IPL and SPL (Cavanna & Trimble, 2006). Functionally, the precunei are part of the semantic network (Binder et al., 2009) but are also strongly associated with episodic memory (e.g., Krause et al., 1999). Given the interrelatedness of semantic and episodic memory, each facilitating information retrieval from the other (Greenberg & Verfaellie, 2010), the similarity of activity in the precunei across the tasks during the period of (presumed) lexical-semantic word planning is consistent with their functional role reported in the literature. Finally, a mainly beta-band-driven similarity was observed in the bilateral lateral occipital areas. While these areas are mainly associated with visual feature processing of objects (e.g., Chouinard et al., 2009; Malach et al., 1995), they might be relevant for language beyond picture naming (e.g., Neudorf et al., 2022). While pre-picture retrieval in context-driven naming is elicited by sentential context, sentence comprehension and subsequent word preplanning might still be accompanied by visual imagery (for haemodynamic results, see Roos et al., 2023). Given that participants are familiarised with the target pictures before the experiment and encounter them twice during the experiment (once per condition), activation of some visual features of the upcoming picture stimuli is plausible.

Across the dissimilar areas, parcels in the frontal lobes and bilateral anterior STG indicated higher power in the alpha and lower beta frequency bands in the object task but higher power in the beta frequency band in the action task. In the right MTG and ITG, however, higher power in the object task was observed not only in the alpha and lower beta frequencies but across the whole range. If we interpret higher alpha power as increased inhibition of an area (Jensen & Mazaheri, 2010; Klimesch et al., 2007), this would suggest that frontal lobes, bilateral anterior STG and right MTG/ITG were more active when retrieving word representations in the action task. Alternatively, and not contradicting the gating-by-inhibition accounts, increased alpha-beta activity could indicate less rich information in the retrieved representations in the object task (Hanslmayr et al., 2012). Both suggestions would be in line with the notion that action naming is more and differentially demanding compared to object naming, amongst other things, due to increased semantic and morphological complexity of verbs (Mätzig et al., 2009). At the neural level, both suggestions are also consistent with the findings that verb processing produces a larger volume of hemodynamic activity compared to noun processing (e.g., Faroqi-Shah et al., 2018).

### Similarity analysis captures coarser lexical-semantic computations in the right hemisphere

We found (dis)similarity predominantly in the right hemisphere (Figure 4C and 4D, Figure S2A-2). While stronger lateralisation of language would be typically expected in the left hemisphere (e.g., Woodhead et al., 2021), some language subprocesses such as sound-to-lexical-meaning mapping (e.g., Bozic et al., 2010) or conceptual retrieval and integration (Binder et al., 2009) have been associated with bilateral language networks. Overall, however, the role of the right hemisphere in language in adults seems to be complementary rather than equipotential to the left hemisphere: When the mature, predominantly left-hemispheric language network is disrupted, the ability of the right hemisphere to contribute to the core language computations remains modest (Wilson & Schneck, 2020). For instance, even when temporo-parietal right-hemispheric areas support word retrieval after left-hemispheric stroke, they are only successful about half of the time (Chupina et al., 2022).

This said, establishing the precise role and the degree of involvement of the right hemisphere in specific language (sub)functions requires further evidence. In their review, Bradshaw and colleagues (2017) reported that the strength and reliability of lateralisation for various language tasks was inconsistent and strongly analysis-driven across studies, depending, amongst other things, on the pre-selected regions of interest and baselines.

Generally, studies using electrophysiological techniques report bilateral activity during word retrieval, including in object and action processing (Levelt et al., 1998; Liljeström et al., 2009; Sörös et al., 2003). In comparison, techniques like fMRI/PET relying on specific data analysis steps such as thresholding commonly report strong left-laterality for language. However, without directly assessing the laterality across homologous ROIs, observing a cluster in the left but not the right hemisphere creates the wrong impression that the activity is left-lateralised whereas, in fact, it is merely stronger in the left hemisphere (see Peelle, 2012).

Martin and colleagues (2022) demonstrated this by showing that the subthreshold BOLD activity was present in right fronto-temporal areas homologous to the left-hemispheric language network nodes engaged in a language comprehension task (see also Turkeltaub et al., 2025).

Analogously to fMRI studies, lack of (dis)similarity in the left-hemispheric posterior language regions in the present study does not, in principle, indicate lack of (dis)similarities across tasks there. Statistically, the fact that we did not find any (dis)similarity in the left temporo-parietal areas merely indicates that the signal between tasks did not become less or more similar when removing this area from the data. The similarity that we found instead in the right hemisphere was interpreted as related to lexical-semantic processes, in line with these areas being homologous to the established context-driven areas in the left hemisphere (e.g., Roos & Piai, 2020). Furthermore, this interpretation is justified by the findings of these right temporo-parietal areas being engaged in the lexical-semantic stages of picture naming measured with MEG (Levelt et al., 1998; Liljeström et al., 2009; Sörös et al., 2003).

Speculatively, our analysis might have highlighted the (dis)similarities in the right hemisphere and masked those in the posterior left hemisphere due to the qualitative computational differences in the bihemispheric language network. In the right hemisphere, higher structural intrahemispheric connectedness has been suggested to be behind higher-correlated activity within the hemisphere and more diffuse electrophysiological responses which might index “coarser” lexical-semantic processing compared to more specialised processing in the left hemisphere (Jung-Beeman, 2005). The neural basis for these distinct computations is likely provided by the microanatomical asymmetries observed across the hemispheres (e.g., Hutsler & Galuske, 2003). Assuming the coarser processing speculation is correct, the fact that this coarse/fine-tuned dichotomy was observed in the posterior but not frontal language areas might indicate the more specialised role of the former, at least in the lexical-semantic processes associated with word retrieval. This suggestion is in line with the literature showing that critical areas for visually guided (for review see Piai & Eikelboom, 2023) as well as context-driven picture naming (Piai et al., 2018) are located in the temporal and parietal regions.

### Degree of abstraction determines conclusions about shared neural processes

Importantly, throughout the paper, we have argued that neural correlates of cognitive activity should be described not only based on their spatial characteristics (i.e., association with brain structures) but also the characteristics of these structures’ neurophysiological responses. Thus, to be able to conclude that two tasks or conditions recruit the same or similar neural mechanisms, we would need to observe the correlates to both overlap spatially *and* in other prespecified neurophysiological parameters.

Following this approach, we defined the similarity between object and action word retrieval tasks as converging temporo-spectral signatures within brain areas. Across the similar areas, this definition allowed for separating activity in the bilateral precuneus from activity in the right parietal-temporal regions based on their differential temporo-spectral profiles, implying different levels of engagement and potentially different functional roles of these areas. Had we defined similarity between object and action word retrieval based solely on spatial overlap, by contrast, we might have concluded that the precunei and parietal areas reflect a shared neural mechanism for object and action words. This illustrates the so-called *abstraction problem* (Francken et al., 2022; Piai et al., 2025): Our conclusions regarding the underlying neural mechanisms are shaped by the degree of abstraction adopted. Although there is no “correct” degree of abstraction for describing a neural process, future investigations should attempt to clearly define which degree of abstraction they assume to be sufficient for claiming (dis)similarity across the compared neural correlates (for discussion see Piai et al., 2025).

### Limitations and future directions

One important limitation of this study concerns the choice of the time window for the similarity analysis across tasks. Identifying the time windows where analogous processes are expected is crucial for a meaningful comparison between tasks or conditions. Here, we chose to align the pre-picture intervals based on an analysis of the trial structure and the observed within-task electrophysiological activity, i.e., the largest decreases of power in the alpha-beta frequency range which are associated with information retrieval (see *Methods* for more detail). While this adjustment of the analysis intervals across the tasks hopefully aligned them better in terms of preplanning processes, it has simultaneously misaligned the intervals in terms of trial timing. At the beginning of the interval, the retrieval signatures in the action task coincided with the auditory presentation of the pronoun *hij* / *zij,* whereas the end of the interval approached picture presentation in the object but not in the action task. While we believe the intervals overall still captured analogous lexical-semantic preplanning processes, we cannot rule out that this realignment had an impact on our results, especially with respect to the time features. This demonstrates that finding optimal temporal alignment between tasks or conditions to capture the same underlying cognitive and neural processes remains challenging and requires careful task design.

Next, while the sentence context in our task was presented in the auditory modality and, thus, word preplanning in the constrained condition was not visually driven, the cognitive processes induced by the picture presentation at the end of the sentence likely impacted the RTs and, potentially, even the pre-picture neural responses. Using a sentence completion task instead, although it might introduce its own challenges (such as higher variability and increased length of responses), could elicit more naturalistic context-driven word production, further disengaging it from visual processes. Finally, to disentangle the potential impact of (micro)saccadic eye movement on the alpha-beta signatures (Popov & Staudigl, 2023) from the task-related language processes, future electrophysiological studies could employ concurrent eyetracking to investigate the mechanisms of the alpha-beta decreases in semantically constraining contexts.

## 5. CONCLUSION

In this study, we aimed to compare neural correlates of object and action word retrieval, focusing specifically on its lexical-semantic stages. To avoid visual confounds across the tasks, typically introduced by visually guided word elicitation, we used context-driven picture naming and analysed the pre-picture retrieval driven by sentential context.

Having defined similarity as converging spectro-temporal signatures within overlapping brain areas, we found that similarity across tasks was linked to right posterior temporal and parietal language areas, whereas dissimilarity was linked to bilateral frontal and anterior-to-medial temporal regions. Importantly, the similarity patterns diverged spectrally for the posterior temporo-parietal region and the precunei, suggesting differential functional mechanisms behind them. We speculatively suggest that lack of similarity in left posterior language areas paired with similarity in right-hemisphere homologous areas might be explained by the degree of computational specialisation of the areas for language. Namely, while the more precise language computations in the left hemisphere might have manifested as smaller (dis)similarities, the coarser representations and/or computations in the right hemisphere could have produced diffuse activity which might have been more clearly (dis)similar in terms of contained mutual information. This speculation would further suggest that the left frontal areas where dissimilarities were found are also less fine-tuned for lexical-semantic stages in word production.

